# The developmental transcriptome of *Parhyale hawaiensis*: microRNAs and mRNAs show different expression dynamics during the maternal-zygotic transition

**DOI:** 10.1101/2021.06.25.449901

**Authors:** Llilians Calvo, Maria Birgaoanu, Tom Pettini, Matthew Ronshaugen, Sam Griffiths-Jones

**Affiliations:** Faculty of Biology, Medicine and Health, The University of Manchester, Manchester, M13 9PT, UK

## Abstract

*Parhyale hawaiensis* has emerged as the crustacean model of choice due to its tractability, ease of imaging, sequenced genome, and development of CRISPR/Cas9 genome editing tools. However, transcriptomic datasets spanning embryonic development are lacking, and there is almost no annotation of non-protein-coding RNAs, including microRNAs. We have sequenced microRNAs, together with mRNAs and long non-coding RNAs, in *Parhyale* using paired size-selected RNA-seq libraries at seven time-points covering important transitions in embryonic development. Focussing on microRNAs, we annotate 175 loci in *Parhyale*, 85 of which have no known homologs. We use these data to annotate the microRNome of 37 crustacean genomes, and suggest a core crustacean microRNA set of around 61 sequence families. We examine the dynamic expression of microRNAs and mRNAs during the maternal-zygotic transition. Our data suggest that zygotic genome activation occurs in two waves in *Parhyale* with microRNAs transcribed almost exclusively in the second wave. Contrary to findings in other arthropods, we do not predict a general role for microRNAs in clearing maternal transcripts. These data significantly expand the available transcriptomics resources for *Parhyale*, and facilitate its use as a model organism for the study of small RNAs in processes ranging from embryonic development to regeneration.

## Introduction

*Parhyale hawaiensis* has emerged as a key crustacean model for studies ranging from regeneration to comparative developmental biology. Available genomics tools include a sequenced genome (1), transcriptome annotation (2,3), and successful application of CRISPR-Cas9 approaches (4). Detailed description of embryonic developmental landmarks such as the segmentation cascade (5), Hox gene expression (6) and cell lineage tracing studies have also been established (7,8). However, there remain a number of missing tools in this expanding repertoire. A key omission is publicly available transcriptome data across the developmental time-course. A few studies using pooled embryos from diverse stages have provided some insight into the *Parhyale* gene and transcript annotation, but there is no genome-wide temporal resolution or information about dynamic expression of transcripts. Existing annotations are limited in sequencing depth and replication, and annotation of small RNAs (including microRNAs) is limited to highly conserved sequences (2,3).

MicroRNAs are short non-coding RNAs of ∼22 nucleotides (nt) in length that regulate gene expression at a post-transcriptional level in metazoans and plants. In animals, microRNAs target the 3’UTRs of mRNAs by partial base-pairing complementarity with target mRNAs (9) inducing either translation inhibition or deadenylation and decay of these target mRNAs (10). Since their discovery, microRNAs have been found to regulate many biological processes, and their importance in development has been demonstrated repeatedly. For example, at the maternal-zygotic transition (MZT), zygotic microRNAs have been found to be involved in clearance of maternally-deposited mRNAs in several invertebrate species including *Drosophila melanogaster* (11) (miR-309 cluster), *Tribolium castaneum* (12) and *Blattella germanica* (also miR-309 cluster) (13). Similar results have been found in vertebrates such as *Danio rerio* (miR-430) (14) and *Xenopus laevis* (miR-427) (15), although the microRNAs involved do not appear to be conserved between vertebrates and invertebrates. Interestingly, this early developmental function for microRNAs has not been found in *Caenorhabditis elegans* (16) or in mice, where it is suggested that microRNAs do not play an essential role in the clearance of maternal mRNAs (17). Indeed, a recent study has shown that only two microRNAs are essential during *C. elegans* development, miR-35 and miR-51 (18). These many studies suggest two principles: first, that microRNAs are key components of embryonic development, either collectively or individually; and second, that variation in microRNA function between species can be significant, and their developmental roles must therefore be examined on a species by species basis.

In this study, we have annotated and quantified the expression of mRNAs, long non-coding RNAs and microRNAs across 7 developmental stages in *P. hawaiensis*. Focusing primarily on microRNA expression during *Parhyale* embryogenesis, we have increased the number of annotated microRNAs in this organism from 51 highly conserved sequences (1,3) to a total of 175 microRNA precursors, 85 of which have not been previously described in any organism. We have used the microRNA repertoire of *Parhyale* to provide a comprehensive homology-based annotation of crustacean microRNAs in 37 species. We find that the core crustacean microRNA complement numbers around 61 families. Finally, the expression dynamics of microRNAs and target mRNAs through development suggests that zygotic genome activation (ZGA) occurs in two waves, with microRNA expression restricted to the second wave.

## Materials and Methods

### *Parhyale hawaiensis* culture, sample preparation and library construction

Wild-type *Parhyale* were kindly donated by Aziz Aboobaker’s lab at Oxford University. Animals were reared in standard plastic aquarium tanks containing artificial sea water (aquarium salt and deionised water) at a salinity of 30 PPT, and kept at ∼26°C. Cultures were aerated with aquarium pumps and airstones, water was changed once a week and animals were fed fish flakes and carrots. Embryos were manually collected from the ventral brood pouch of gravid females anaesthetised using clove oil (Sigma) diluted 1:10000 in sea water. Embryos were washed in filtered seawater and manually staged using a Leica Stereo Fluorescence microscope. Isolated embryos were stored in RNAlater (Sigma) and total RNA was then extracted using the SPLIT RNA Extraction Kit (Lexogen) following the manufacturer’s instructions. Small RNA libraries (4 replicates per time-point) were built using the Small RNA-Seq Library Prep Kit (Lexogen). Long library fragments and linker-linker artefacts were removed using a purification module with magnetic beads (Lexogen). Long mRNA libraries (2 replicates per time-point) were built using the TruSeq Stranded mRNA HT Sample Prep Kit (Illumina). Library concentration was assessed for all libraries using the Qubit fluorimetric system (Invitrogen) and quality was assessed using the Agilent 2200 TapeStation. Sequencing was performed at the University of Manchester Genomic Technologies Facility.

### Small RNA-seq data analysis and microRNA prediction

RNA-seq raw reads were trimmed using Cutadapt v. 1.18 (19) and read length distribution was assessed using FastQC v0.11.8 (20). For microRNA predictions, reads ranging from 18 to 26 nucleotides (nt) were retained. These reads were mapped to *Parhyale* tRNAs and rRNAs using Bowtie (v1.1.1; parameters -p4 -v3 --un) (21). *Parhyale* tRNAs were predicted using tRNAscan-SE (v2.0; option -e) (22,23) and crustacean rRNAs were downloaded from RNAcentral release 16 (24). Non tRNA/rRNA reads were then mapped to the *P. hawaiensis* genome (Phaw 5.0; GCA_001587735.2) allowing only one mismatch (Bowtie; -v 1 -S -a –best –strata), and the mapped reads were used for microRNA annotation using the miRDeep2 tool (v 0.1.1) (25). To run Mirdeep2 we used all the metazoan microRNAs available on miRBase as references (v 22.1) (26). Predicted microRNAs were manually filtered to keep microRNAs obeying the following criteria (27): at least 10 reads for both the 5p and 3p mature sequences, minimum loop length of 8 nt, and at least 50% of the reads for each mature microRNA having the same 5’ end. Exceptions were only made for highly conserved microRNAs that are confidently annotated in other species, predicted using BLASTN (v2.6.0+; -word_size 4 - reward 2 -penalty -3 -evalue 0.01 -perc_identity 70) (28) hits and verified by manual inspection.

### Identification of *Parhyale* microRNA homologs in crustacean genomes

We downloaded the genomes of 37 crustacean species available in NCBI (Table 1) and searched for homologs of all the 175 predicted microRNAs in *Parhyale* using BLASTN (-word_size 4 -evalue 0.01 -reward 2 -penalty -3 -perc_identity 30). A presence/absence matrix of microRNA families and family copy number was plotted in R using the package pheatmap (v1.0.12) (29).

**Table 1:** Crustacean genomes from NCBI

| Specie | NCBI Assembly |
| --- | --- |
| *Hyalella azteca* | GCA_000764305.2 Hazt_2.0 |
| *Triops cancriformis* | GCA_000981345.1 tcf_1.0 |
| *Caridina multidentata* | GCA_002091895.1 Cmul_gen_Assembly01 |
| *Ligia exotica* | GCA_002091915.1 Lexo_gen_Assembly01 |
| *Penaeus japonicus* | GCA_002291165.1 Mjap_WGS_v1 |
| *Procambarus virginalis* | GCA_002838885.1 Pvir0.4 |
| *Eulimnadia texana* | GCA_002872375.1 clam_shrimp_assembly_v0.1 |
| *Eriocheir sinensis* | GCA_013436485.1 ASM1343648v1 |
| *Lepidurus apus* | GCA_003723985.1 Lubb2018 |
| *Lepidurus arcticus* | GCA_003724045.1 ASM372404v1 |
| *Penaeus vannamei* | GCA_003789085.1 ASM378908v1 |
| *Daphnia magna* | GCA_003990815.1 ASM399081v1 |
| *Palaemon carinicauda* | GCA_004011675.1 ASM401167v1 |
| *Armadillidium vulgare* | GCA_004104545.1 Arma_vul_BF2787 |
| *Pandalus platyceros* | GCA_005815305.1 GSC_Sprawn_1.0 |
| *Trinorchestia longiramus* | GCA_006783055.1 ASM678305v1 |
| *Tigriopus californicus* | GCA_007210705.1 Tcal_SD_v2.1 |
| *Penaeus monodon* | GCA_015228065.1 NSTDA_Pmon_1 |
| *Portunus trituberculatus* | GCA_008373055.1 ASM837305v1 |
| *Armadillidium nasatum* | GCA_009176605.1 CNRS_Arma_nasa_1.0 |
| *Cherax quadricarinatus* | GCA_009761615.1 DU_Cquad_1.0 |
| *Amphibalanus amphitrite* | GCA_009805615.1 SNU_Aamp_1 |
| *Cherax destructor* | GCA_009830355.1 DU_Cdes_1.0 |
| *Pollicipes pollicipes* | GCA_011947565.2 Ppol_2 |
| *Tigriopus kingsejongensis* | GCA_012959195.1 ASM1295919v1 |
| *Caligus rogercresseyi* | GCA_013387185.1 ASM1338718v1 |
| *Daphnia dubia* | GCA_013387435.1 dubia_v0.01 |
| *Platorchestia sp* | GCA_014220935.1 ASM1422093v1 |
| *Semibalanus balanoides* | GCA_014673585.1 Sbal3.1 |
| *Orchestia grillus* | GCA_014899125.1 Ogril_1 |
| *Macrobrachium nipponense* | GCA_015104395.1 ASM1510439v1 |
| *Daphnia pulex* | GCA_900092285.2 PA42_4.1 |
| *Oithona nana* | GCA_900157175.1 O_NANA_1 |
| *Acartia tonsa* | GCA_900241095.1 Aton1.0 |
| *Apocyclops royi* | GCA_900607525.1 AroyWGS1.0 |
| *Tisbe holothuriae* | GCA_900659605.1 ASM90065960v1 |
| *Eurytemora affinis* | GCF_000591075.1 Eaff_2.0 |

### Relative arm usage

Homologs of *Parhyale* precursors in three other species with expression data available – *Tribolium castaneum, Apis mellifera* and *Parasteatoda tepidariorum* – were identified by BLASTN (-task blastn-short -evalue 0.01) and manual inspection. Read counts for *T. castaneum* mature microRNAs were obtained from Ninova et al. (12), counts for *A. mellifera* from Pires et al. (30) and counts for *P. tepidariorum* were calculated in-house using methods and data from Leite et al. (31). Relative arm usage was calculated using the method described in Marco et al. (32): log2(N5’/N3’); where N5’ is the number of reads mapped to the -5p arm, and N3’ is the number of reads mapped to the -3p arm.

### Transcriptome assembly and annotation

Paired-end RNA-seq reads from each developmental library were mapped to the *Parhyale* genome (Phaw 5.0; GCA_001587735.2) using STAR (v2.7.2b) (33). The mapped reads were then assembled using Trinity (v2.9.0) (34). and the resulting transcripts were mapped back to the genome with gmap (version 2020-06-01) (35), and duplicates removed. Only these transcripts were used for further analysis. Transdecoder (v5.5.0) (https://github.com/TransDecoder/TransDecoder) was used to identify potential coding regions within these transcripts and only the longest ORF per transcript was kept. Using BLAST search (Uniprot release 2020_02, BLASTP version 2.9.0, e-value ≤ 10^−6^), we looked for ORFs with homology to known proteins in 7 crustacean species with annotated transcriptomes (*Daphnia pulex, Daphnia magna, Penaeus vannamei, Armadillidium nasatum, Armadillidium vulgare, Portunus trituberculatus*, and *Tigriopus californicus*), as well as *Drosophila melanogaster* and *Apis mellifera*.

### Functional annotation

We searched for protein signatures in the Pfam database (version 33.1) (36) using hmmscan (HMMER version 3.3.2) (37). Using BLAST (BLASTP version 2.9.0; e-value ≤ 10^−3^), we searched our peptide sequences against annotated Swissprot proteins (Uniprot release 2021_02). The Pfam and Uniprot hits were then loaded into a Trinotate sqlite database (v3.2.2) (38), which provided KEGG, EGGNOG and GO terms associated with each transcript.

### Quantification and differential gene expression analysis

To quantify small RNAs, reads not mapping to tRNAs/rRNAs were mapped to the predicted mature microRNAs, using Bowtie (v1.1.1; -v 1 -S -a) and mapped reads were quantified using salmon quant from salmon (v0.14.1) (39) in alignment mode. To quantify mRNAs, reads mapped to the annotated transcriptome were quantified using salmon quant from salmon (v0.14.1) in mapping base mode.

Quantifications were then used for differential expression analysis using the package DESeq2 (v1.28.1) (40) in R Studio (v1.3.1056) (41) R v4.0.2 (42). To group each mRNA or microRNA into expression clusters we applied fuzzy c-means clustering using the function cmeans from the R package e1071 (v1.7.4) (43) to the normalized TPM computed using DESeq2. The number of clusters for each dataset was previously determined using the elbow method. Heatmaps were computed using the R package ComplexHeatmap (v2.5.5) (44). The proportion of previously annotated vs newly annotated microRNAs belonging to each expression cluster was assessed by chi-squared tests, using the ratio within the total population of expressed microRNAs to generate expected values.

### Target prediction

Targets of our annotated microRNAs within the predicted UTRs of our annotated mRNA transcripts were predicted using Seedvicious (v1.3) (45) and filtered adhering to the following criteria: free energy below -10 Kcal/mol, microRNA expressed with at least 10 RPM, mRNA expressed at least 10 TPM, each microRNA targets the same UTR more than once. Pairwise Mann-Whitney-Wilcoxon tests were performed between all possible pairs of the five different mRNA expression clusters, to compare the number of different microRNAs targeting each mRNA and the number of microRNA targeting sites per mRNA 3’UTR. Enrichment of microRNA-targeted mRNAs in each mRNA cluster was assessed using the phyper formula of the hypergeometric distribution in R as follows: phyper(q-1, m, n, k, lower.tail = FALSE), where q = successes in subset, m = successes in population, n = population total - successes in population, and k = subset.

### Gene Ontology (GO) annotation and GO enrichment analysis

Significantly upregulated mRNAs were subjected to gene set enrichment analysis using the TopGO package v2.42.0 in R (46). The classic Fisher test was used to generate enrichment p-values, with the algorithm weight01 and a p-value cutoff of p< 0.01.

### Data Access

All RNA sequencing data and quantifications were deposited in the Gene Expression Omnibus (GEO) database under accession number GSE178877.

## Results

### *Parhyale hawaiensis* size-separated RNA sequencing and small RNA annotation

To develop a comprehensive developmental transcriptome of *Parhyale*, we selected embryos from seven different key time-points spanning the whole of embryogenesis. The time-points were chosen to capture key transcriptional changes during important developmental transitions (Figure 1A). The first time-point covers the 1 to 8 cell stages (S1-4) – at this time the zygote is still transcriptionally inactive (47), and therefore has exclusively maternally-loaded RNAs. The second time-point contains the 32-cell stage, named stage 6 (S6), described in the literature as the maternal-to-zygotic transition (47). During the third time-point, stages 8 to 11 (S8-11), embryonic cells migrate and segregate from the yolk cells. The next two time-points were built using precisely staged embryos at stage 14 (S14) and stage 17 (S17) during the period of germ band extension. The final two libraries span stages 21 to 23 (S21-23) and 24 to 30 (S24-30), during which limb buds form and morphogenesis occurs (Figure 1A). To facilitate comparison of microRNA and mRNA expression profiles during embryogenesis, we built paired “small” (<150nt) and “large” (>150nt) libraries from the same samples for each time-point (Figure 1B).

**Figure 1.**
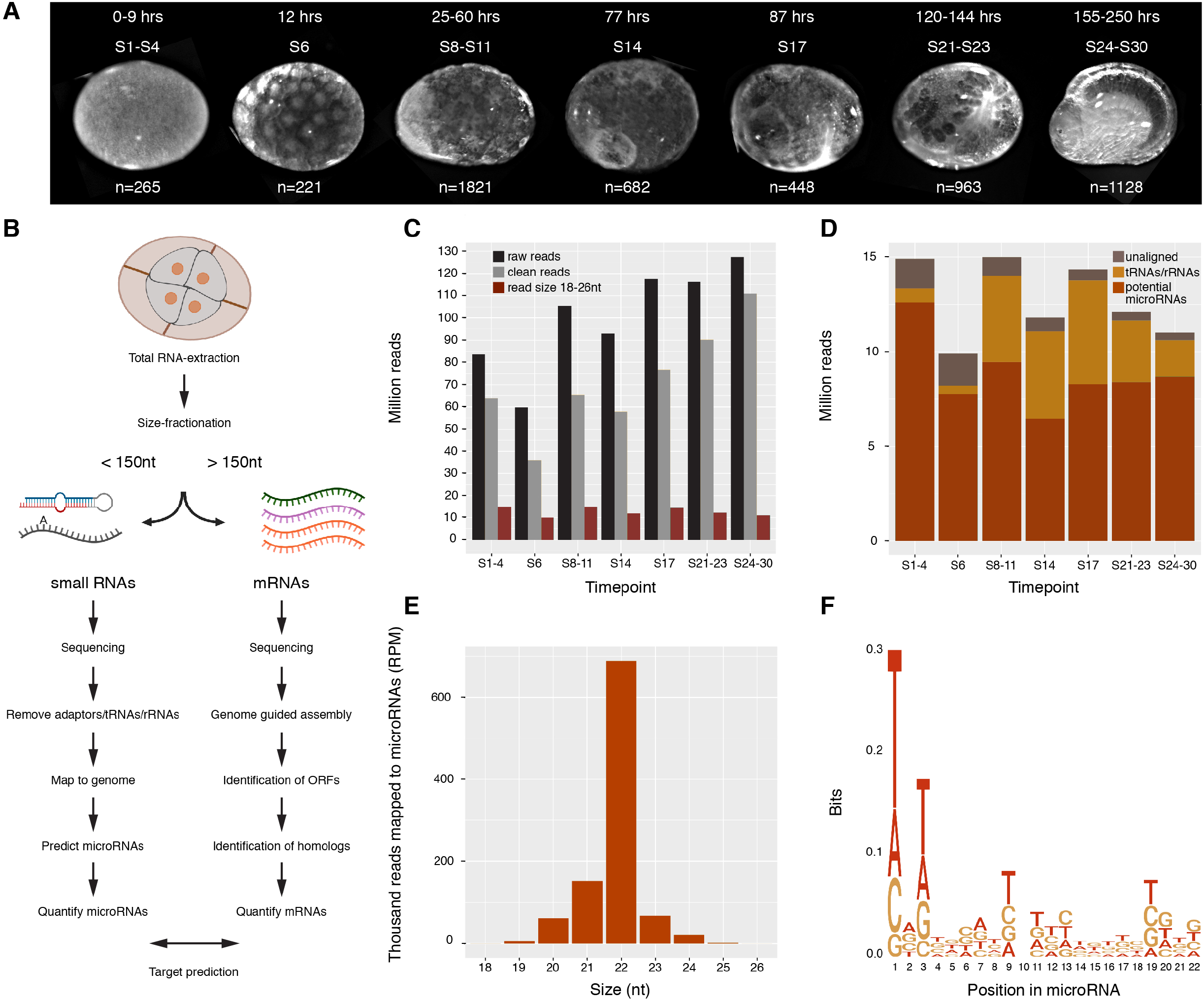
Library preparation and microRNA annotation in *Parhyale* development. (**A**) Brightfield images of embryo stages selected for building libraries. Developmental stages, the corresponding number of hours post-fertilization, and the number of embryos used for each time-point are indicated (**B**) Diagram of workflow for size-separated library preparation and analysis. (**C**) Absolute abundance of sequence reads per time-point, total sequences reads (black), clean reads remaining after adaptor removal (grey), reads remaining after size selection (brown). (**D**) Distribution of size selected reads following mapping to the genome and to tRNAs/rRNAs database. (**E**) Size distribution of reads mapping to predicted microRNAs. (**F**) Sequence-logo of the first 22 nt of non-redundant reads mapping to microRNAs.

The small RNA reads obtained from the sequencing were cleaned (adaptors removed) and selected to retain 18-26nt reads (Figure 1C). Reads that mapped to the genome but failed to map to *Parhyale* tRNAs or crustacean rRNAs were considered potential microRNAs and used for miRDeep2 microRNA prediction (Figure 1D). Manual inspection of miRDeep2 predictions yielded a total of 175 high confidence microRNA precursors, and 349 distinct mature sequences. 87 of the precursor loci were conserved among other metazoans, and 88 were previously unreported. As expected, the majority of the reads mapping to the predicted microRNAs were 22nt long (Figure 1E) and 5’ uracil biased (Figure 1F).

### Annotation of predicted *Parhyale* microRNAs in crustacean genomes

MicroRNAs in crustaceans are poorly annotated. Only *Daphnia pulex, Marsupenaeus japonicus* and *Triops cancriformis* have any published microRNA sequences, and the level of coverage and completeness is variable. In order to address this underlying sampling problem, we used the 175 microRNA precursors identified from our sequencing data in *Parhyale* to predict microRNA homologs in the genomes of 37 crustacean species available in NCBI. In *Parhyale*, the 175 identified precursors belong to 105 different microRNA families; 51 families have been previously annotated, and are therefore conserved with other metazoans, whereas 54 families were novel. 124 out of 175 precursors, belonging to 79 different families, were present in the genome of at least one other crustacean species surveyed (see Figure 2), with 18 families not conserved outside of the Malacostraca. We therefore suggest that the core crustacean microRNA set is comprised of around 61 families. A total of 49 out of 175 precursors, belonging to 26 of the novel families, had no significant match in any other crustacean genome, and are therefore ‘*Parhyale* unique’ (28% of all precursor sequences). The 37 crustacean species tested include four species in the same order as *Parhyale* (Amphipoda). Divergence times among these five amphipod species is not well determined, but all belong to the Talitroid clade, sharing a common ancestor ∼60 million years ago, therefore indicating that these 49 ‘*Parhyale* unique’ precursors have evolved more recently than ∼60 million years ago (48).

**Figure 2.**
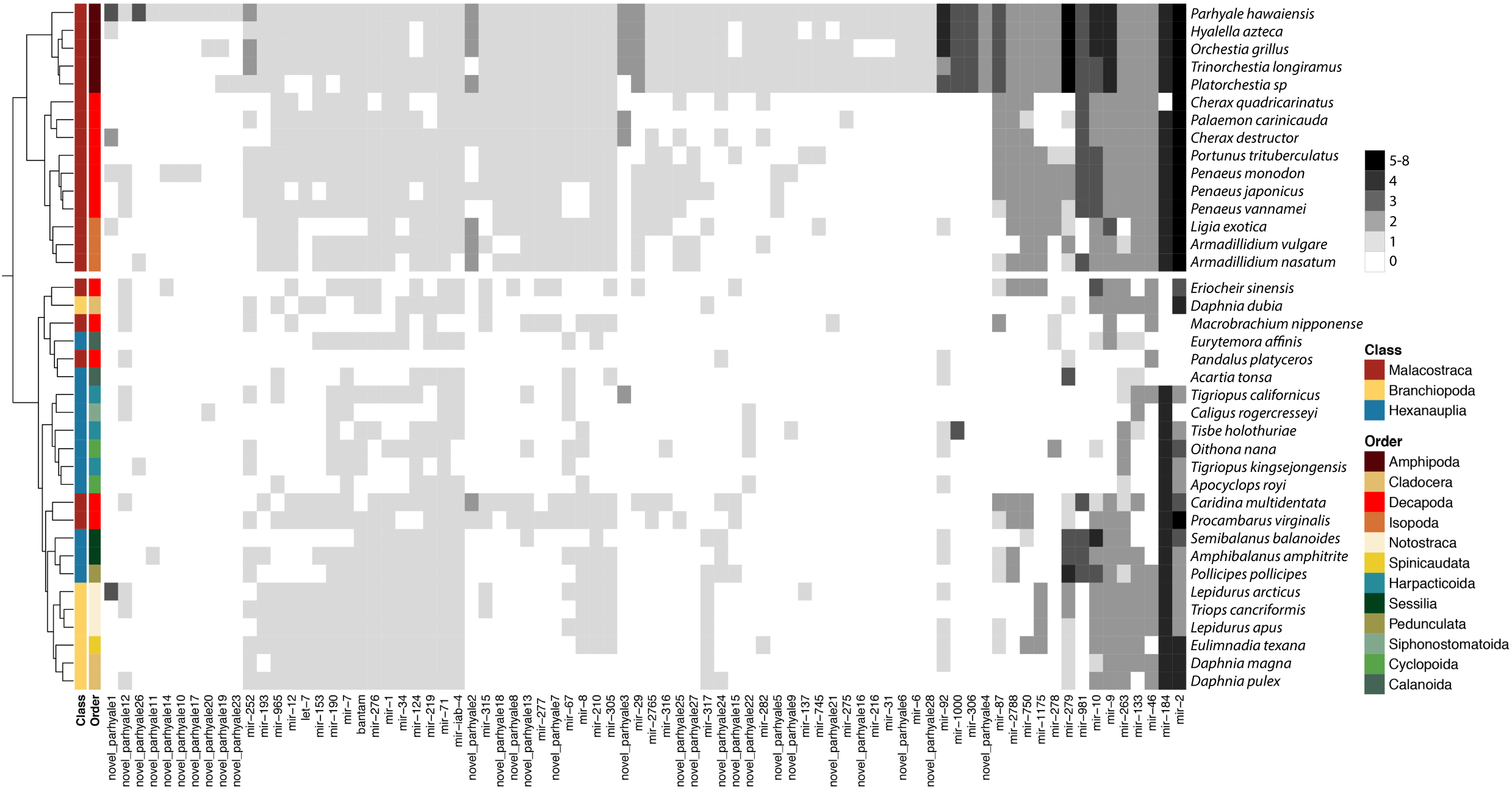
Homologs of *Parhyale* microRNAs in annotated crustacean genomes from NCBI. Heatmap representing *Parhyale* microRNA families in the genomes of 37 crustaceans; greyscale indicates the number of members per microRNA family found in each species. Clustering analysis based on microRNA presence/absence was used to generate a species tree (left). The class and order of each species is indicated by the colour-coded ribbon.

We clustered the set of crustacean species based on the presence and absence of microRNA families in their genomes. The resulting tree closely reproduces aspects of the existing established phylogeny of the crustacea (Figure 2). For example, *Parhyale* has more microRNAs in common with other members of the class Malacostraca than it does with species belonging to the more distant Branchiopoda and Hexanauplia classes. Similarly, we observe strong correspondence in microRNA presence among species within the same order as *Parhyale*, the Amphipoda.

### *Parhyale* microRNA arm switching in development and evolution

Each microRNA hairpin precursor is processed to produce two possible mature products, often of unequal abundance. Historically, the less abundant product was termed the miR* sequence, and was presumed to be degraded. Recently, this view has been abandoned with the discovery that for some animal microRNAs, arm dominance can differ between tissues, developmental times or species. Additionally, studies have shown that each arm can have many different targets, and that both arms can be functional. Arm switching therefore has the potential to diversify microRNA function (49,50).

We have examined developmental and evolutionary arm switching of all the predicted *Parhyale* microRNAs (Figure 3A). Almost all microRNAs showed the same dominant arm throughout the course of development. However, a small proportion of microRNAs exhibit developmental arm switching (Figure 3A). For some microRNAs, a pronounced switch in dominance was observed across the short timescale of adjacent time-points, for example LQNS02278186.1_47223 and LQNS02277275.1_26288. For many microRNAs, approximately equal proportions of 5p and 3p arms were detected at specific time-points (Figure 3A, white tiles), suggesting that both potential sequences may function to target different mRNAs at the same stages (49).

**Figure 3.**
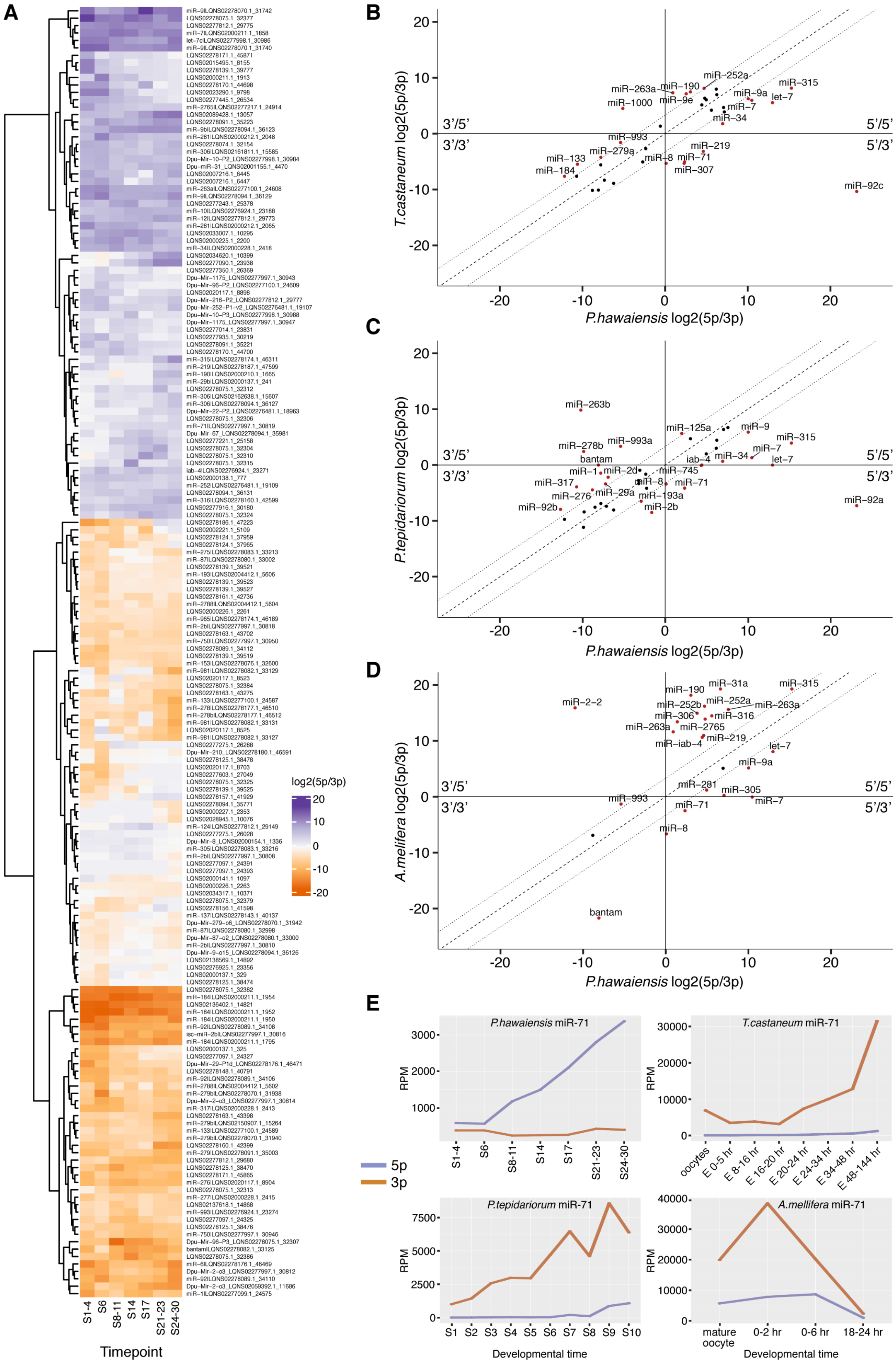
MicroRNA arm switching through development and evolution. (**A**) Heatmap showing relative arm usage changes across seven time-points of *Parhyale* embryonic development. Purple indicates 5’ dominance, orange 3’ dominance. **(B)** Comparison of the relative arm usage of microRNA homologs between *Parhyale* and *Tribolium* **(B)**, spider **(C)** and honeybee **(D)**. All three show microRNAs that have undergone arm switching (5’/3’ or 3’/5’ quadrants). Dotted lines show the 10-fold difference boundary, name labels are shown for each microRNA exceeding this 10-fold change in arm usage. (**E**) miR-71-5p and -3p arm expression data through development for the four species analysed.

By comparing arm dominance between datasets for different species, we also identified some cases of arm switching through evolution (Figure 3B-D). For example, miR-71 is consistently 3’-biased in the flour beetle, spider and honeybee (Figure 3B, C, D respectively), but 5’-biased in *Parhyale* (Figure 3E), suggesting that miR-71 switched arms in evolution during arthropod diversification. miR-8 is also switched in *Parhyale* when compared to the other three species, although less dramatically, changing from a 3’ bias in all three species, to approximately equal arm usage in *Parhyale* (Figure 3B, C, D).

### MicroRNA expression dynamics in embryogenesis

We have used normalised read counts to quantify the expression of all 349 predicted *Parhyale* mature microRNAs throughout embryogenesis. Principle component analysis (PCA – Figure 4A) using the mature read counts confirmed high similarity among replicates within each time-point, and also showed high similarity between the first two time-points (S1-4 and S6). These findings were confirmed with Spearman correlation tests for all pairwise combinations of expression profiles among the seven time-points (Figure 4B). Previous estimates of zygotic genome activation place the timing between S4 and S6, and therefore large-scale expression differences might be expected between our first two time-points. Our observation of similar profiles between S1-4 and S6 suggests that zygotic transcription of microRNAs has not yet begun at S6. In contrast, the S8-11 time-point is clearly separated from the earlier stages in the PCA analysis, and the correlation coefficient between S6 and S8-11 is the lowest of any pair of adjacent time-points (Figure 4B). We therefore suggest that the onset of zygotic microRNA expression occurs between S6 and S8. The mid-stages of embryogenesis (S8-11, S14, S17) show a similar microRNA composition (r>0.9), which is distinct from the early stages. The microRNA composition by the end of embryogenesis (S24-30) is markedly different from both early and mid-embryogenesis (r<0.7). The penultimate time-point (S21-23) is transitionary between mid and late embryogenesis.

**Figure 4.**
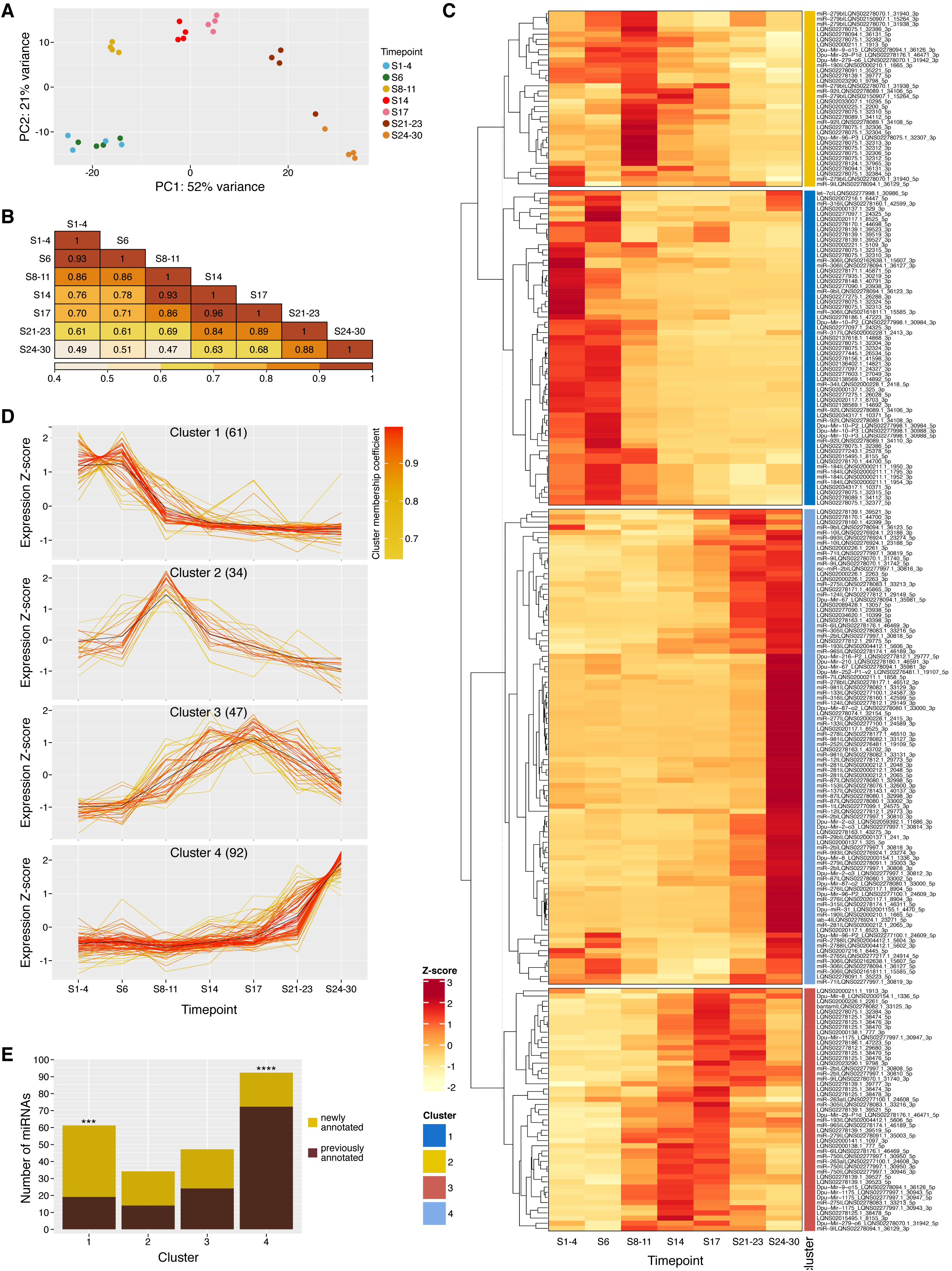
Differential expression analysis of microRNAs during development. (**A**) Principal component analysis (PCA) of microRNA expression in each replicate and time-point. Four replicates are shown for each time-point. (**B**) Heatmap of all-versus-all pairwise Spearman correlation coefficients between time-points. Numbers in tiles are r values, and heatmap colour coding is based on r value. **(C)** Heatmap showing z-score calculated for expression of each microRNA through embryonic development. Each microRNA is classified into an expression cluster, indicated by the colour coded ribbon. **(D)** Expression profiles for microRNAs with membership scores ≥ 0.6 for each cluster; the number indicated in parentheses is the total number of microRNAs belonging to each cluster. (**E**) Composition of the 4 expression clusters in newly annotated and previously annotated microRNAs. Chi-squared tests were performed for each cluster. Cluster 1 shows a significantly higher proportion of newly annotated microRNAs than expected, *X*^2^ (1, *N* = 61) = 14.95, p = 1.10⨯10^−4^, whereas cluster 4 contains an unexpectedly high proportion of conserved microRNAs, *X*^2^ (1, *N* = 92) = 19.40, p = 1.05⨯10^−5^, p-value significance levels are indicated by asterisks.

Of the 349 mature microRNAs annotated, 234 (67%) had relatively high expression levels (≥ 10 RPM) in at least 1 time-point (see Figure 4C). We find that at least 172 mature microRNAs are maternally provided in the fertilized egg (S1-4), whereas the number of expressed microRNAs drops slightly to 167 by S6. This drop is likely due to degradation without additional microRNA transcription, consistent with the suggestion that zygotic transcription of microRNAs does not occur until after S6. The number of expressed microRNAs is steady throughout mid-embryogenesis (S8-S11: 163; S14: 164; S17: 165), with a slight increase in the last two time-points (S21-23: 174; S24-30: 178).

MicroRNAs with similar expression profiles across the time course are likely to be involved in similar developmental processes. We therefore used a fuzzy c-means clustering approach to group microRNAs with similar expression dynamics. Unlike k-means, fuzzy c-means clustering assigns a membership coefficient to each microRNA, such that each data point belongs to a greater or lesser degree to each cluster. Using this approach with the 234 mature microRNAs highly expressed in at least one stage, we identified four expression clusters (Figure 4C). Expression profiles of the most significant microRNAs for each cluster (membership cut-off 0.6) are shown in Figure 4D. Cluster 1 (26% of the 234 mature microRNAs) comprises exclusively maternally loaded microRNAs, which are expected to function in the early embryo even before the onset of ZGA. Cluster 2 is composed of the first microRNAs to be expressed during ZGA (15%), cluster 3 represents microRNAs expressed predominantly during mid embryogenesis (20%), and cluster 4 includes the microRNAs expressed almost exclusively at late embryogenesis (39%) (Figure 4D). We see that the largest number of microRNAs are expressed in late embryogenesis. This finding is similar to results reported in other species including zebrafish (51), *Drosophila virilis* (52) and *Tribolium* (12) where more microRNAs were found to be expressed at later stages, but different from findings in mice (53). The high number of microRNAs with peak expression in later stages correlates with the increase in the number and variety of differentiated cell-types.

Comparing the distribution of conserved versus newly annotated microRNAs in each cluster revealed that cluster 1 (expressed during early embryogenesis) contains a disproportionately high number of newly annotated microRNAs (42 new, 19 conserved; *X*^2^ (1, *N* = 61) = 14.95, p = 1.10⨯10^−4^), whereas cluster 4 (expressed late in embryogenesis) contains a significantly higher proportion of conserved microRNAs (20 new: 72 conserved, *X*^2^ (1, *N* = 92) = 19.40, p = 1.05×10^−5^) (Figure 4E). This abundance of young or novel microRNAs in early stages has also been described in other arthropod species such as *Drosophila virilis* (52), *Tribolium* (12) and *Blatella germanica* (54).

### mRNA expression dynamics in embryogenesis

To compare mRNA and small RNA expression dynamics in *Parhyale*, we use Trinity-based pipeline to perform genome guided annotation of the developmental transcriptome using poly-A selected RNA-seq datasets collected across the same samples as above. We annotated a total of 49,532 protein-coding transcripts from 31,087 Trinity genes. Details of the transcriptome annotation pipeline and statistics are shown in Table 2.

**Table 2:** Transcriptome statistics

| **Transcriptome statistics** |  |
| --- | --- |
| Total trinity genes | 31087 |
| Total trinity transcripts | 49532 |
| Percent GC | 46.46 |
| Contig N10 | 10065 |
| Contig N20 | 7591 |
| Contig N30 | 6090 |
| Contig N40 | 4918 |
| Contig N50 | 3934 |
| Average length (bp) | 2146.44 |
| Median length (bp) | 1211.5 |
| Total assembled bases | 106317470 |
| **3'UTRs statistics** |  |
| Transcripts with 3'UTRs | 42505 |
| Average length (bp) | 1135.70 |
| Median length (bp) | 319 |
| Max length (bp) | 22970 |
| Min length (bp) | 1 |
| Sequences >10nt | 30427 |

PCA analysis of transcript expression profiles defined by read counts clearly shows good agreement between the two replicates per time-point (Figure 5A). As with the microRNA analysis, both PCA analysis and Spearman’s correlation coefficients (Figure 5B) show that the mRNA content at the two earliest stages of embryogenesis (S1-4 and S6) is similar, but markedly different from later time-points. Correlation scores show that the final stage (S24-30) is also very different from all preceding time-points, indicating that a distinct set of mRNAs is engaged at the end of embryogenesis, presumably in establishing the final RNA profiles of adult tissues (Figure 5B).

**Figure 5.**
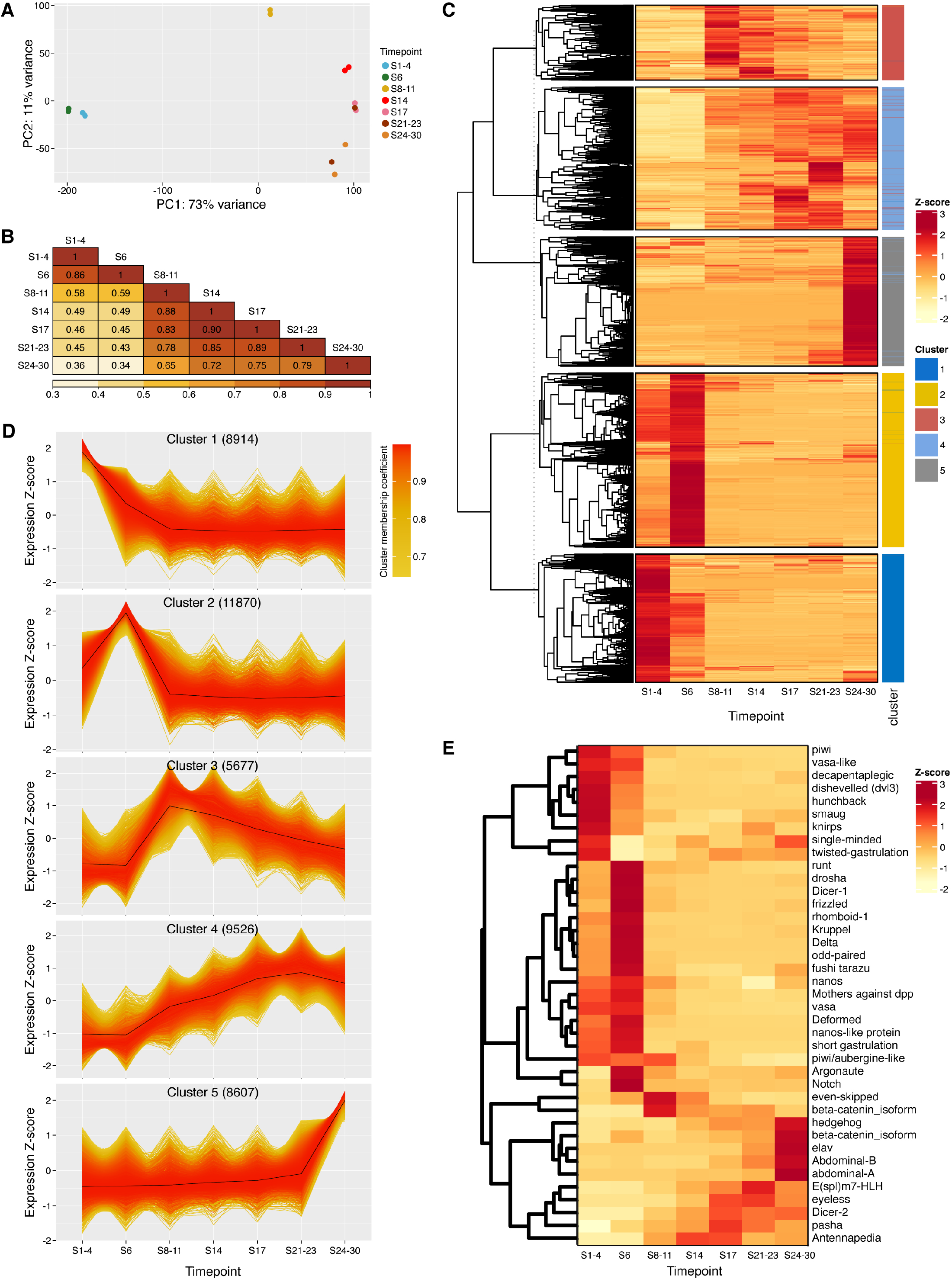
Differential expression analysis of mRNAs during development. (**A**) Principal component analysis (PCA) of mRNA expression levels in each replicate and time-point. Two replicates are shown for each time-point. (**B**) Heatmap of all-versus-all pairwise Spearman correlation coefficients calculated between time-points. Numbers in tiles are r values, and heatmap colour coding is based on r value. **(C)** Heatmap showing z-score calculated for expression of each mRNA through embryonic development. Each mRNA is classified into an expression cluster, indicated by the colour coded ribbon. **(D)** Expression profiles for mRNAs with membership scores ≥ 0.6 for each cluster; the number indicated in parentheses is the total number of mRNAs belonging to each cluster. (**E)** Heatmap showing z-score of a subset of known developmental genes extracted from (D).

A total of 44,594 mRNAs had relatively high expression levels (≥ 10 TPM) in at least one time-point (Figure 5C). As with microRNAs, we clustered mRNA expression profiles using fuzzy c-means clustering yielding five different clusters. Expression profiles of the most significant mRNAs for each cluster (membership cut-off 0.6) are shown in Figure 5D. Cluster 2 represents mRNAs with peak expression at S6 (Figure 5C and D), immediately after the time of ZGA previously reported in *Parhyale* (47). This cluster is entirely absent from the microRNAs expression profiles (Figure 4C and D), suggesting that microRNAs are not generally transcribed during the first stages of ZGA. Indeed, 47% of the mRNA transcripts belong to clusters 1 and 2, representing peak expression at the first two time-points, whereas only 26% of microRNAs belong to the cluster that includes high expression at the first two time-points. Conversely, only 19% of mRNAs fall into cluster 5 (peak expression at S24-30), whereas the equivalent cluster 4 for microRNAs contained 39% of the microRNAs. These data clearly suggest that zygotic expression of many mRNAs is initiated before zygotic microRNAs are expressed, and that mRNAs have functional roles early in embryogenesis independent of microRNAs, while a high proportion of microRNAs function late.

We examined the expression levels of a number of specific genes known to play important roles during embryogenesis or in microRNA biogenesis in other species (Figure 5E). For example, mRNAs including *nanos, hunchback, dishevelled* and *smaug* are all known to be maternally loaded in other species, and predicted homologs were also present at high level in the *Parhyale* early embryo. In *Drosophila*, the RNA-binding protein Smaug is an important player during MZT, responsible for the degradation of hundreds of maternally loaded mRNAs (55); in *Parhyale, smaug* homolog expression is highest at S1-4, consistent with a conserved biological function.

In accordance with other studies in *Parhyale*, our analysis failed to detect a clear *zelda* ortholog. However, we found that a predicted homolog of *odd-paired*, a pioneering factor suggested to be a key player during ZGA (56), was maternally loaded and its expression increased at S6, showing the same behaviour as *Dmel zld*. Expression of homologs of other *Drosophila* pair-rule genes (*eve, ftz* and *runt*) also increased during ZGA. Conservation of temporal expression was also observed for mRNAs known to be expressed during late embryogenesis, such as *eyeless, elav* and *E(spl)m7-HLH* involved in eye development, axon guidance and neurogenesis respectively. Expression of *piwi* and *vasa*, components of the piRNA processing pathway are predominantly expressed in the early embryo, whereas *Dicer-2* (implicated in siRNA processing) is primarily expressed at mid to late stages. This hints that piRNAs are likely to play important roles in the early embryo, whereas siRNA function may be more prominent later in development. Interestingly, of the proteins required for microRNA processing, *Dicer-1* and *drosha* mRNA expression peak early at S6 with sustained expression thereafter, whereas *pasha* expression peaks around mid to late embryogenesis.

### Comparative expression dynamics of microRNAs and their predicted targets

Our paired, size separated libraries allow the analysis of temporal expression of microRNAs in combination with their targets (mRNAs). Target predictions were performed using the SeedVicious algorithm and potential interactions then filtered (see methods) to produce a list of putative microRNA-mRNA interactions. To assess the degree to which mRNAs are targeted by microRNAs through development, we analysed the proportion of mRNAs in each expression cluster that are predicted to be targeted by microRNAs (Figure 6A). Of the total 44,594 mRNAs assigned to expression clusters, 28,129 (63%) had 3’ UTRs, and of these, 6,883 (15.4% of total mRNAs) were predicted to be targeted by microRNAs. Using a hypergeometric test, we find that clusters 1 and 4 were significantly under-enriched for mRNAs targeted by microRNAs, whereas cluster 2 was significantly over-enriched for targeted mRNAs (Figure 6A). Cluster 1 primarily contains maternally loaded mRNAs, whereas cluster 2 represents the first wave of zygotic transcripts.

**Figure 6.**
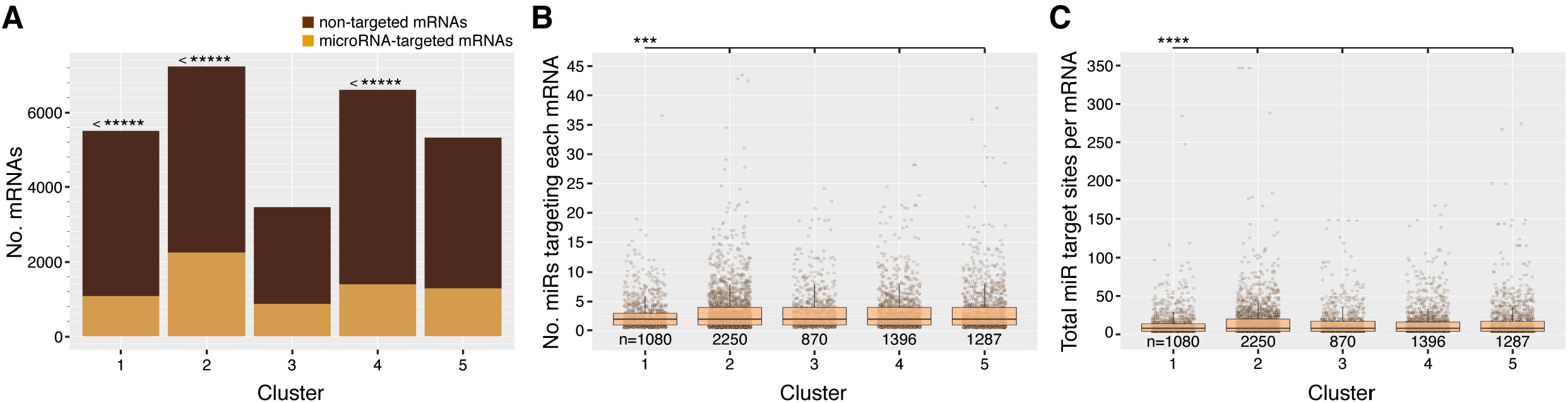
Differential targeting of mRNAs by microRNAs through development. (**A**) The number of mRNAs in each expression cluster that are predicted to be targeted by microRNAs versus those that are non-targeted. Hypergeometric tests were performed to compare the observed numbers with expected values, calculated based on the overall proportion of targeted mRNAs within the entire mRNA population. Significance values are indicated by asterisks. Cluster 1 is under-enriched for targeted mRNAs, p= 1.21⨯10^−21^, cluster 2 is over-enriched for targeted mRNAs, p= 1.02⨯10^−50^, and cluster 4 is under-enriched for targeted mRNAs, p= 1.34⨯10^−13^. **(B)** The number of different microRNAs targeting each mRNA in each expression cluster. Pairwise Mann-Whitney-Wilcoxon tests were performed between all clusters. Cluster 1 mRNAs are targeted by significantly fewer microRNAs than mRNAs in all other clusters. Cluster 1 vs 2, p= 4.66⨯10^−6^, cluster 1 vs 3, p= 7.12⨯10^−6^, cluster 1 vs 4, p= 1.46×10^−4^, cluster 1 vs 5, p= 2.79⨯10^−4^. All other pairwise comparisons p>0.05. **(C)** The number of microRNA targeting sites per mRNA 3’UTR in each expression cluster. Pairwise Mann-Whitney-Wilcoxon tests were performed between all clusters. Cluster 1 mRNAs have significantly fewer microRNA target sites than mRNAs in all other clusters. Cluster 1 vs 2, p= 5.48⨯10^−7^, cluster 1 vs 3, p= 8.67⨯10^−6^, cluster 1 vs 4, p= 6.93⨯10^−6^, cluster 1 vs 5, p= 1.02⨯10^−5^. All other pairwise comparisons p>0.05.

We also compared the number of different microRNAs targeting each mRNA per expression cluster (Figure 6B), and the number of microRNA target sites in each 3’ UTR per cluster (Figure 6C). Pairwise Mann-Whitney-Wilcoxon tests between the five clusters in all combinations revealed that mRNAs in cluster 1 were targeted by significantly fewer different microRNAs than all other clusters, and also had significantly fewer microRNA target sites in their 3’ UTRs than mRNAs in all other clusters. These data therefore suggest that globally, microRNAs are more involved in regulating zygotically expressed genes than in clearing maternally loaded transcripts.

### Differential expression analysis of mRNAs and microRNAs identifies two waves of ZGA

To further explore the apparent developmental lag in zygotic expression of microRNAs with respect to mRNAs, we compared the expression levels of mRNAs and microRNAs between adjacent time-points, S1-4 with S6, and S6 with S8, using DESeq2 (Figure 7). Changes between S1-4 and S6 could be due to degradation of maternally loaded RNAs, or onset of zygotic RNA production. We find that the expression of only one microRNA (LQNS02278075.1_32386_3p) increased significantly (log2 fold change >1.5, padj <0.05) between S1-4 and S6 (Figure 7A). The overwhelming majority of microRNAs either decreased, likely signifying degradation without replacement, or did not significantly change. In contrast, a total of 41 different microRNAs were significantly upregulated between S6 and S8-11, representing 17.5% of all microRNAs expressed during development (Figure 7B). An equivalent analysis of the mRNAs showed that 3568 mRNAs (8% of all expressed mRNAs in development) were significantly upregulated (log2 fold change >1.5, padj <0.05) between S1-4 and S6 (Figure 7C). Between S6 and S8-11, a total of 10288 mRNAs (23.1%) were upregulated (log2 fold change >1.5, padj <0.05) (Figure 7D). We therefore propose a model where a subset of protein-coding genes are activated in a first wave of ZGA between S1-4 and S6, with expression of microRNAs accompanying a larger number of mRNAs that are activated at a later point during the second (and biggest) wave of ZGA between S6 and S8-11 (Figure 7E). Additional equivalent analysis of putative lncRNAs revealed a pattern of activation intermediate between microRNAs and mRNAs, with 3,665 lncRNAs (1.3% of all expressed lncRNAs) significantly upregulated (log2 fold change >1.5, padj <0.05) between S1-4 and S6, and 16,560 (6.0%) significantly upregulated between S6 and S8-11 (Supplementary Figure 1). Gene set enrichment analysis suggests that mRNA transcripts upregulated in the first wave are enriched for Gene Ontology terms related to metabolic pathways, whereas mRNAs upregulated during the second wave are enriched for RNA binding activity and translation (Supplementary Figure 2).

**Figure 7.**
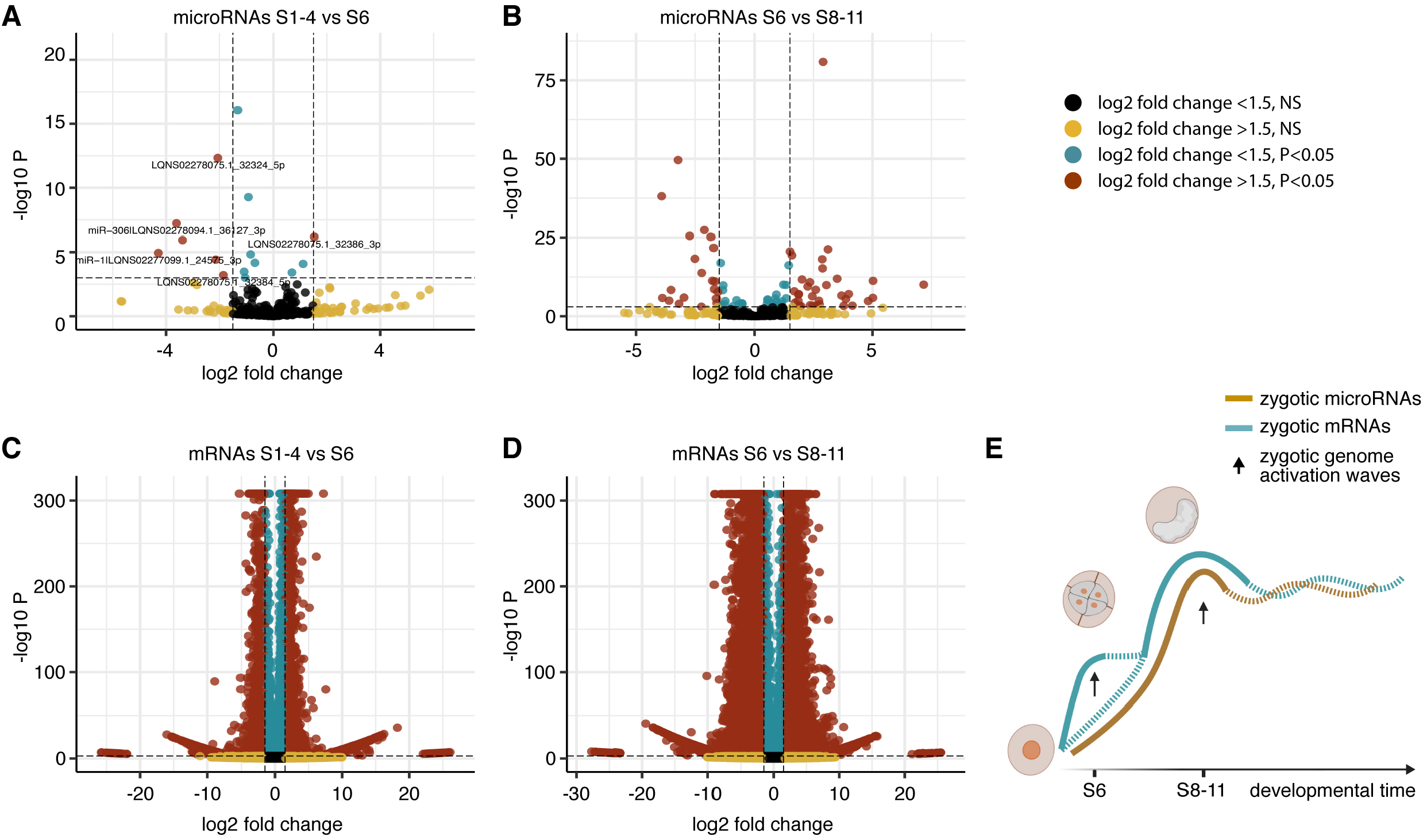
Differential expression analysis during zygotic genome activation. **(A** and **B)** Volcano plots showing log2 fold change in expression (x-axis) versus the p-value (y-axis), for each microRNA expressed between the first two time-points S1-4 to S6 (A) and S6 to S8-11 (B). (**C** and **D**) Volcano plot showing log2 fold change in expression (x-axis) versus the p-value (y-axis) for each mRNA expressed between the first two time-points S1-4 to S6 (C) and S6 to S8-11 (D). Only red dots (log2 fold change ≤ -1.5 or ≥ 1.5 with padj < 0.05 are considered significant. (**E**) Model of zygotic genome activation occurring in two different waves of expression for the mRNAs; onset of microRNA expression occurs only in the second wave.

## Discussion

We have generated paired, size-separated libraries across different stages of *Parhyale* embryogenesis, providing both microRNA and mRNA developmental transcriptomes for this crustacean model organism. We have identified a total of 349 mature microRNAs expressed from 175 microRNA precursors in the genome of *Parhyale*. Of the precursor loci, 87 have been previously described while 88 are unrelated to any previously identified microRNAs in any species. We have used this dataset to provide a first glimpse at the microRNome of 37 other crustacean species, the majority of which have no microRNA expression data or annotation available. This work therefore enables and accelerates investigation of crustacean microRNAs.

Analysis of microRNA expression shows that the early embryo is a highly permissive environment, over-enriched for evolutionarily younger microRNAs, an observation that we can now extend from the Holometabolan insects to the Pancrustaceans (52). This emerging theme suggests insights into microRNA birth, selection, and death processes. We also find that very conserved microRNAs dominate the later stages of development, in agreement with previous studies (12,52).

Our data suggests that ZGA in *Parhyale* occurs in two waves. In the first wave, transcription is primarily associated with a set of early mRNAs and lncRNAs, while the onset of microRNA transcription is almost exclusively limited to the second wave, along with additional mRNAs and lncRNAs. This observation of two waves of ZGA has been reported in several other species (57). In *Drosophila*, the first wave of transcription is widespread across the genome, producing short, inefficiently processed transcripts (58,59). The function of transcription at these loci could be to activate regions of the genome for later transcription of competent mRNAs. However, many of the short early transcripts have been found to be implicated in the sex-determination pathway, thus suggesting function beyond genome activation (59). Furthermore, specific sets of genes are also known to be expressed in two waves in mice; for example, paternal genes are expressed preferentially during the minor ZGA. In the chicken, the opposite was observed with the paternal transcriptome only being activated during the second wave (60). Interestingly, also in chicken, microRNAs are predominantly transcribed during the second wave of ZGA, as we see in *Parhyale* (60).

Target prediction showed that maternally-loaded mRNAs are targeted by fewer microRNAs and have fewer microRNA target sites than later expressed mRNAs. We also see that a large proportion of microRNAs show peak expression at the very end of embryogenesis, suggesting that in *Parhyale*, microRNAs might be more active players during the late stages of development. This is in contrast to the mRNAs, almost half of which show peak expression at the earliest two time-points, while a relatively small proportion peak at late embryogenesis. These findings all point to microRNA regulation being more prevalent for zygotic genes than for maternally-loaded transcripts. This may reflect the importance of microRNAs in balancing and buffering active transcription (61), a role less necessary for maternal transcripts that are not being actively replenished, and which may instead degrade with time via other passive or active mechanisms.

For decades, *Drosophila* has held a virtual monopoly over transcriptomics studies in arthropods, and much of our knowledge today about development is thanks to the fly community. However, other organisms provide models for evolutionary questions that cannot be tackled in *Drosophila* alone. We provide annotation of the *Parhyale* transcriptome (both mRNAs and microRNAs) throughout embryonic development. To our knowledge this is the first publicly-available study in *Parhyale* that provides temporal resolution throughout embryogenesis with tightly spaced time-points, and representation of major developmental transitions such as ZGA, germ band extension, and morphogenesis. This work helps to establish *Parhyale* as a model for questions related to the evolution of crustaceans and insects and facilitates functional studies of microRNAs during crustacean development.

## Acknowledgements

We thank Aziz Aboobaker and his lab for kindly providing starter cultures of *Parhyale* and training. We are grateful to the University of Manchester Genomic Technologies facility, particularly Beverley Anderson, Claire Morrisroe, and Andy Hayes for sequencing and assistance. We thank Hilary Ashe for support and discussions, and the Wellcome Trust for supporting this work via a PhD studentship (203990/Z/16/A) to L.C.

## Author contribution statement

Conceptualization L.C., M.R. and S.G.J.; culture and embryo handling L.C. and T.P.; RNA-Seq L.C.; data analysis L.C.; microRNome annotation L.C.; transcriptome annotation M.B.; statistical analysis L.C. and T.P.; writing – draft L.C.; writing – review and editing L.C., T.P., M.R., and S.G.J.; funding from the Wellcome Trust – PhD studentship to L.C.

## Supplementary figures legends

**Supplementary figure 1.**
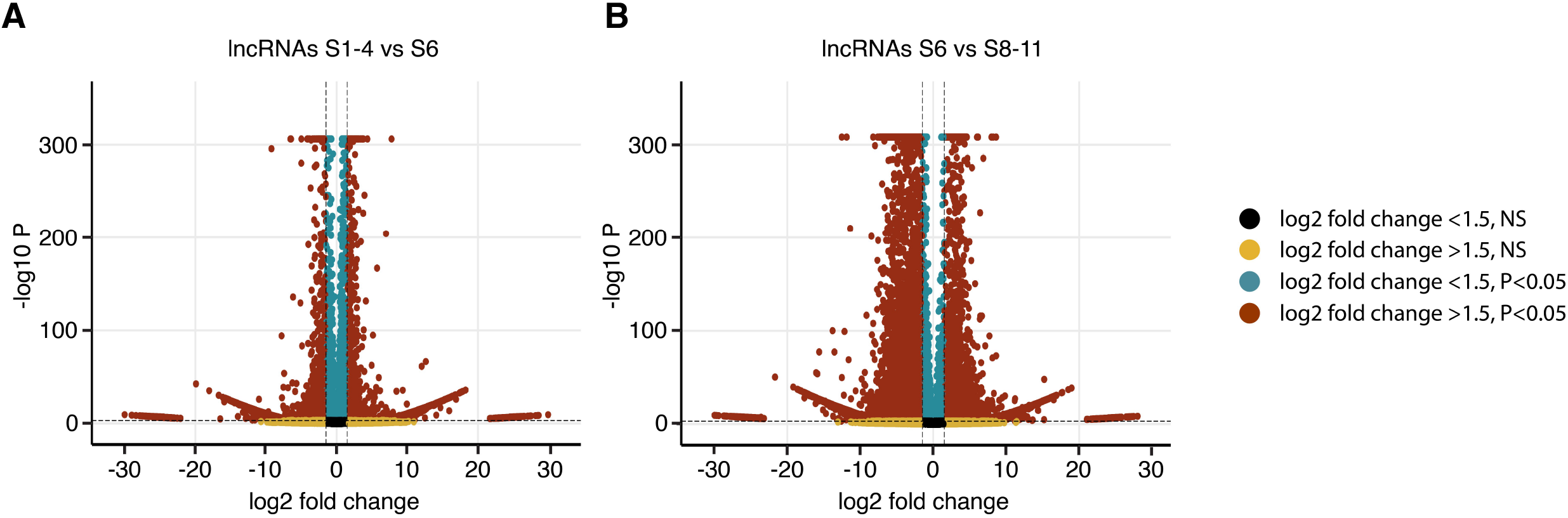
Differential expression analysis of lncRNAs during zygotic genome activation. **(A** and **B)** Volcano plots showing log2 fold change in expression (x-axis) versus the p-adjusted value (y-axis), for each lncRNA expressed between the first two time-points S1-4 to S6 (A) and S6 to S8-11 (B). Only red dots (log2 fold change ≤ -1.5 or ≥ 1.5 with padj < 0.05 are considered significant.

**Supplementary figure 2.**
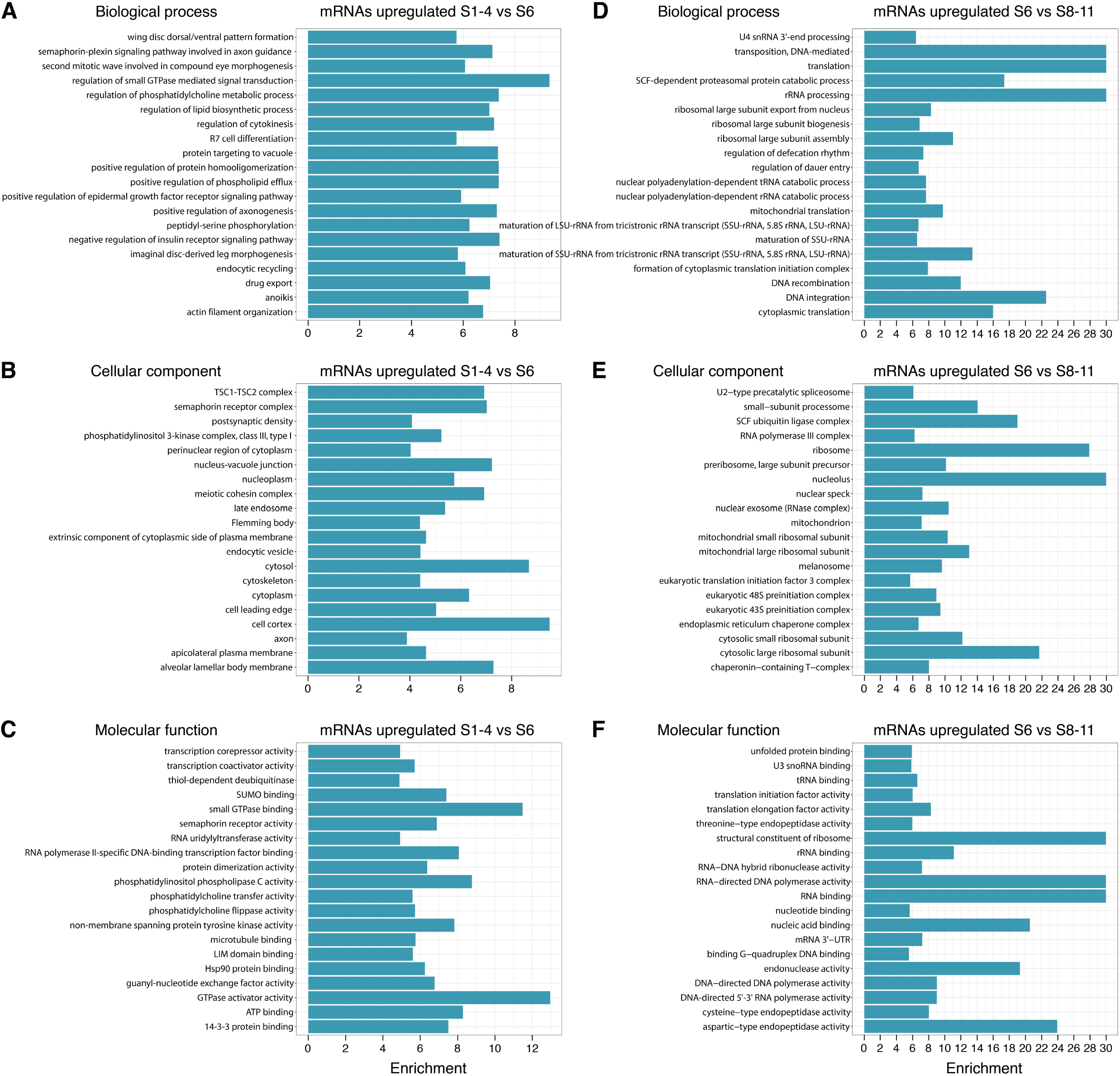
TopGO enrichment analysis results of upregulated mRNAs during the two waves of zygotic genome activation. Functional enrichment analysis was performed for the upregulated mRNAs between S1-4 and S6 **(A, B, C)** and between S6 and S8-S11 **(D, C, F)**. The top 20 enriched GO terms belonging to GO Biological Processes, GO Cellular Components, and GO Molecular Functions are shown for each comparison respectively.

## Notes

### Competing Interest Statement

The authors have declared no competing interest.

